# MMM: A deterministic Mendelian mismatch matrix framework for germplasm collection quality assessment, demonstrated in apple and sweet cherry

**DOI:** 10.64898/2026.08.02.742285

**Authors:** Qiliang Chen, Liuxiu Chen, Jing Fan, Xiaoping Yang, Jingguo Zhang, Wei Du, Wei Liu, Zi’ang Liu, Hongju Hu

## Abstract

Pedigree errors, duplicate accessions, and undocumented kinship limit the effective use of large genebank collections. Existing tools such as CERVUS and COLONY require allele-frequency estimates and return relationship-level assignments, providing no collection-wide view of Mendelian consistency. We developed the Mendelian Mismatch Matrix (MMM) framework and its open-source implementation, MMMAS (Mendelian Mismatch Matrix Analysis System). We applied MMM to apple (*Malus × domestica*; 1,085 accessions, 15 SSRs) and sweet cherry (*Prunus avium*; 383 accessions, 13 SSRs) datasets from the German Fruit Genebank. MMM converts pairwise Mendelian mismatch numbers (Mmn) into an N × N symmetric matrix, from which we derive four collection-scale indices (Mmin, the Mendelian minimum mismatch number; Mavg, the Mendelian average mismatch number; Mzmp, the Mendelian zero-mismatch partner number; Mmmd, the Mendelian mismatch-mode duplication index) and the Mendelian Exhaustive Stratification (MES) grading system. Apple showed a gradient-type structure (Mendelian Grade-gap Index, MGI = 0.125), sweet cherry a broken-type structure (MGI = 0.545). The Mmin–Mavg strategy identified 20 and 10 distinct accessions in apple and sweet cherry, including the scab-resistance donor *Malus × floribunda* 821 (Rvi6) and the fire-blight donor *M. × robusta* 5. The Mmmd index flagged one literature-confirmed pair of duplicate genotypes in sweet cherry. Of CERVUS-confirmed parent–offspring pairs, 97.31% in apple and 100% of high-confidence pairs in sweet cherry fell within Mmn ≤ 1. Mzmp rankings matched documented breeding history, with ‘Cox Orange’ (Mzmp = 57) and ‘Lübecker Bunte’ (Mzmp = 49) ranking first in each species. The complete end-to-end analysis of all 588,070 apple pairwise comparisons took 7.24 s on an ordinary workstation. This deterministic pre-screening framework, with a parameter-free core, supports routine quality assessment in germplasm collections.

## 1. Introduction

Germplasm collections underpin breeding and genetics research (Brown 1989) and have expanded steadily worldwide (Anglin *et al*. 2025; Li *et al*. 2026). In China, national fruit-tree repositories built over seven decades now hold more than 30,000 fruit accessions across multiple institutions (Liang *et al*. 2025). Declining genotyping costs have accelerated this expansion (Anglin *et al*. 2025), so curators and breeders now face practical decisions: separating duplicates from unique accessions, identifying reliable parents, and deciding which genotypes to prioritize for conservation and crossing. Apple (*Malus ×domestica* Borkh.) and sweet cherry (*Prunus avium* L.) illustrate these challenges: long generation times and decades of historical breeding have produced complex pedigree networks (Peace 2017; Muranty *et al*. 2020; Reim *et al*. 2023), even as chromosome-scale reference genomes such as the telomere-to-telomere assembly of sweet cherry ‘Tieton’ continue to expand the molecular resources available for these species (Yu *et al*. 2025a).

Molecular markers have become standard for parentage verification and genetic diversity assessment in horticultural crops, a priority reflected in a recent journal editorial on germplasm and molecular breeding (Wu *et al*. 2023). Simple sequence repeat (SSR) markers have been applied to both genetic diversity assessment and parentage verification, the latter demonstrated in Japanese pear (Sawamura *et al*. 2008), apple (Broschewitz *et al*. 2024), and sweet cherry (Reim *et al*. 2023), while diversity and structure have been characterized in pear (Ouni *et al*. 2020) and across seventeen Chinese native Malus species (Gao *et al*. 2021); they remain widely used in national genebanks because of their lower cost, broader accessibility, and established protocols (Anglin *et al*. 2025). Beyond parentage assignment, molecular assays also support routine genotype certification, such as variety fingerprinting in sweetpotato (Meng *et al*. 2018) and S-RNase genotype determination in Malus (Liu *et al*. 2024). Single nucleotide polymorphism (SNP) arrays and genotyping-by-sequencing now offer higher throughput for research-grade pedigree reconstruction (Vanderzande *et al*. 2019; Muranty *et al*. 2020). Existing parentage programs (Kalinowski *et al*. 2007; Huisman 2017; Cockburn *et al*. 2021), relatedness estimators (Queller and Goodnight 1989; Manichaikul *et al*. 2010), and core-collection algorithms (Kim *et al*. 2007; De Beukelaer *et al*. 2018; Nie *et al*. 2021; Tao *et al*. 2023) operate at the pairwise or subset scale. None offers a deterministic, parameter-free, collection-wide synthesis based on exact Mendelian exclusion. Genotyping errors (typically 0.1%–1% per locus in SSR data) are often neglected in curation workflows (Pompanon *et al*. 2005; Hoffman and Amos 2005). Large germplasm collections also lack a deterministic pre-screening layer that maps conflicts and redundancy before likelihood-based refinement and that remains interpretable when error rates are unknown (Anglin *et al*. 2025). Mendelian exclusion is not new: it has long been applied to parentage testing and data quality control for individual candidate pairs (Jamieson and Taylor 1997; Blouin 2003; Arias *et al*. 2022). Its structural use, however, differs fundamentally between prior work and the present framework. Muranty *et al*. (2020) applied Mendelian error counting as a pairwise exclusion test: a scalar mismatch count per postulated parent–offspring pair was compared against an empirical threshold and then discarded after the accept/reject decision, so no collection-level metrics were retained. In contrast, MMMAS (Mendelian Mismatch Matrix Analysis System) retains the complete N × N mismatch matrix as its primary analytical substrate. This supports row-wise, column-wise, and whole-matrix operations that pairwise-only frameworks cannot define. The pairwise-exclusion workflow of Muranty *et al*. (2020) thus serves as a terminal refinement tool when candidate relationships are already hypothesized, whereas MMMAS operates without prior relationship assumptions and produces a collection-wide diagnostic map.

To address this gap in germplasm management workflows, we developed the Mendelian Mismatch Matrix (MMM) framework, implemented as the open-source software MMMAS. The framework converts the pairwise Mendelian mismatch number (Mmn) into an N × N symmetric matrix and derives collection-scale diagnostics of genetic position, connectivity, and duplication, together with a Mendelian stratification grading system. The framework applies to any diploid species; here we demonstrate it in apple and sweet cherry, and discuss broader applicability in Section 4. All diagnostics are computed in a single pass without allele-frequency estimates or iterative optimization, making the core computation deterministic and parameter-free even when genotyping error rates are unknown; only the downstream exploratory thresholds (e.g., the quadrant cut-offs in functional stratification) are dataset-dependent.

The objectives of this study were to (1) demonstrate the utility of deterministic pre-screening for routine genebank quality assessment using apple and sweet cherry SSR datasets from the German Fruit Genebank, (2) validate mismatch-based parentage detection against published CERVUS assignments, (3) identify genetically distinct accessions and breeding hubs for conservation prioritization, and (4) establish minimum marker-panel requirements and benchmark computational scalability on medium-to-large collections.

## 2. Materials and methods

### 2.1. Datasets

#### Apple

Raw genotypic data were obtained from the OpenAgrar public repository of the German Fruit Genebank (Julius Kühn-Institut) (Broschewitz *et al*. 2023): 1,404 apple (*Malus ×domestica* Borkh.) accessions genotyped at 17 nuclear SSR markers recommended by the European Cooperative Programme for Plant Genetic Resources (ECPGR) *Malus/Pyrus* Working Group. Samples annotated as triploid or tetraploid were removed, as were samples displaying three or more alleles at any locus (mixed samples or cryptic polyploidy), leaving 1,085 diploid accessions for Mendelian exclusion analysis, which assumes diploid segregation (Blouin 2003). Null allele frequencies were estimated with CERVUS 3.0.7 (Kalinowski *et al*. 2007), a maximum-likelihood estimator whose accuracy has been evaluated by simulation (Chapuis and Estoup 2007); two loci, CH05e03 (F(Null) = 0.157) and CH04f10 (F(Null) = 0.223), were excluded under the criterion F(Null) > 10% to reduce false-positive mismatches caused by null alleles (Dakin and Avise 2004). The final panel comprised 15 SSR loci with a mean of 20.0 alleles per locus, a mean polymorphism information content (PIC) of 0.811, and a cumulative two-parent exclusion probability of > 99.9999%.

#### Sweet cherry

Sweet cherry genotypic data were likewise obtained from OpenAgrar (Höfer *et al*. 2021): 383 diploid accessions genotyped at 16 SSR markers. All accessions were diploid; no polyploid samples required removal. Under the same criterion, three loci (PS05C03, F(Null) = 0.224; PceGA34, F(Null) = 0.194; UDP98-412, F(Null) = 0.143) were excluded. The final panel comprised 13 loci with a mean PIC of 0.589 and a cumulative two-parent exclusion probability of 99.9997%.

### 2.2. The MMM framework

#### Compatibility assessment

The Mendelian exclusion logic underlying the MMM framework builds on established theoretical foundations for pedigree verification. Khan *et al*. (2017) evaluated rule-based procedures for resolving Mendelian inconsistencies in nuclear pedigrees through simulations of 20,000 nuclear families typed for two-allele markers, showing that the detectability of genotype errors depends on marker polymorphism and family role, and proposing a set of improved data-cleaning rules. Calus *et al*. (2011) showed that opposing homozygotes (pairs of individuals sharing no alleles at a locus) provide effective indicators for detecting Mendelian inconsistencies between SNP genotype data and recorded pedigree relationships; Arias *et al*. (2022) further classified Mendelian errors in SNP-array data and separated genomic alterations from calling errors. These principles transfer directly to horticultural germplasm, where codominant SSR markers are routinely applied.

The MMM framework applies to diploid codominant markers: SSR, SNP and insertion/deletion (InDel) markers. The empirical thresholds and validation reported here are based on SSR data, and SNP-specific considerations are addressed in Section 4.5. For a population of N diploid individuals scored at L loci, the genotype of individual i at locus l is the unordered allele set *G_i,l_={a^(1)^_i,l_, a^(2)^_i,l_}*. The Boolean mismatch function is

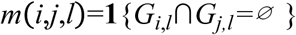

where **1**{⋅} is the indicator function: when two individuals share no allele at a locus, a parent-offspring relationship is excluded at that locus by Mendel’s law. The Mendelian mismatch number (Mmn) of pair (i, j) is the count of incompatible loci:

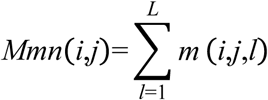

satisfying 0 ≤ Mmn(i,j) ≤ L, Mmn(i,j) = Mmn(j,i) and Mmn(i,i) ≡ 0. The number of informative loci L(i, j) is the count of loci with non-missing genotypes in both individuals, and the mismatch rate r(i, j) = Mmn(i, j)/L(i, j) expresses genetic difference as a proportion. All N(N − 1)/2 pairwise evaluations form a symmetric integer matrix **M**, with time complexity O(N²L) and space complexity O(N²).

#### Individual-level indices

From **M**, we compute three indices that jointly characterize an individual’s genetic position within the collection. The Mendelian minimum mismatch number (Mmin(i)=min{j≠iMmn(i,j) measures proximity to the nearest neighbor: a small value implies that at least one other accession is genetically close. The Mendelian average mismatch number (Mavg(i)=1/N−1Σ_j≠1_ M mn(i,j)) captures distinctness from the entire collection.

A complementary statistic, the Mendelian zero-mismatch partner number (Mzmp(i)=|{≠iMmn(i,j)=0}|), quantifies local connectivity density. High-Mzmp accessions are typically historical core breeding parents or widely used pollinizers. Mmin and Mzmp are coupled at the binary level: Mzmp = 0 whenever Mmin ≥ 1, and Mzmp ≥ 1 whenever Mmin = 0. Mmin thus marks only whether an accession has any zero-mismatch partner, whereas within the Mmin = 0 subset Mzmp resolves how many such partners it has.

Duplicate detection uses the Mendelian mismatch-mode duplication index (Mmmd): two accessions are flagged as duplicates when their mismatch vectors against all third-party accessions are element-wise identical over loci genotyped in both. This criterion has no pairwise analogue: direct pairwise Mmn = 0 can also arise from full-sib or parent–offspring relationships. Allele dosage and homozygous/heterozygous state are intentionally discarded, trading relationship-category resolution for parameter independence and computational efficiency.

#### Functional quadrant classification

Mzmp counts zero-mismatch partners but does not reveal whether they derive from a narrow local pedigree or from divergent lineages; Mavg, the mean mismatch against the entire collection, supplies that missing dimension. Accessions were therefore classified by the joint Mavg–Mzmp distribution into four quadrants using three dataset-dependent exploratory thresholds: (i) a connectivity cutoff placed in the upper tail of the Mzmp distribution (capturing approximately the top 3% of accessions); (ii) the median of the Mavg distribution; and (iii) the Tukey upper fence of the Mavg distribution (Q3 + 1.5 × IQR, where IQR is the interquartile range; Tukey 1977). Accessions above the connectivity cutoff and at or above the median were classed as bridge accessions, those above the cutoff but below the median as pedigree cores, those below the cutoff but at or above the Tukey fence as genetic extremes, and all remaining accessions as standard. The dataset-specific values of these thresholds are reported in Section 3.5.

#### Mendelian Exhaustive Stratification (MES)

Accessions are partitioned by Mmin into mutually exclusive and exhaustive difference grades *D_k_*={*i:Mmin(i)=k*} and nested cumulative grades *C_k_*={*i:Mmin(i)*≥*k*}, with *C_k_*_+1_⊆*C_k_*and *C*_0_ equal to the entire collection; D₄⁺ denotes grade D₄ and above. The Mendelian Grade-gap Index (MGI) 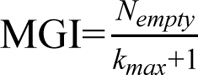, where *N_empty_* is the number of empty grades between 0 and the maximum observed grade *k_max_*, quantifies structural continuity: low values indicate a gradient-type structure, high values a broken-type structure.

MES grades are sample-dependent: they express relative distinctness under the current collection composition and marker panel, not an intrinsic cross-study property. Unlike core-collection tools that maximize allelic diversity (PowerCore, Kim *et al*. 2007; Core Hunter, De Beukelaer *et al*. 2018), MES stratifies the existing collection by internal genetic structure without removing any accession.

### 2.3. Marker-panel sensitivity analysis

Seven nested SSR configurations were constructed from the apple dataset (N = 1,085): the original 17 SSRs, the 15 SSRs remaining after null-allele filtering, and panels of 13, 11, 9, 7 and 5 loci by sequentially removing the lowest-PIC loci (mean PIC = 0.819, 0.811, 0.827, 0.837, 0.847, 0.853 and 0.860, respectively). Loci were removed sequentially from lowest to highest PIC to provide a conservative estimate of the minimum panel size required to maintain diagnostic resolution, mimicking practical optimization constraints in which low-PIC loci are the first candidates for exclusion during panel downsizing. Evaluation metrics were the MES grade distribution, the highest attainable grade, the number of D₄⁺ accessions, and the grade stability of tracked accessions.

### 2.4. Validation design

#### Pedigree concordance

Apple results were compared with published CERVUS parentage assessments at the 95% logarithm of odds (LOD) threshold, covering 253 cultivars and 381 putative parent–offspring pairs, of which 335 were CERVUS-confirmed and 46 were CERVUS-rejected (Broschewitz *et al*. 2024). Sweet cherry results were compared with the independent assessment of Reim *et al*. (2023): 56 cultivars with 112 putative relationships, of which 87 parent–offspring pairs had genotypes for both members and were stratified by trio/pair confidence (*, 95%; +, 80%; −, < 80%). Taking the published CERVUS assignments as the reference, concordance was summarized by two ratios: the proportion of CERVUS-confirmed pairs falling within Mmn ≤ 1, and the proportion of Mmn ≤ 1 candidates confirmed by CERVUS at the 95% LOD threshold (Kalinowski *et al*. 2007). The strict Mmn ≤ 1 criterion was chosen because parent–offspring pairs sharing no alleles at two or more loci are incompatible with Mendelian inheritance under the assumed error-free model, consistent with the exclusion thresholds applied in SSR-based parentage studies (Jamieson and Taylor 1997; Dakin and Avise 2004).

#### Distinctness and hub identification

Distinct accessions were identified along two paths: path A selected accessions with Mmin ≥ 4 (D₄⁺) to capture local distinctness, and path B applied Tukey’s fences to the collection-wide Mavg distribution (threshold Q3 + 1.5 × IQR; Tukey 1977) to capture global distinctness. Hubs were identified by Mzmp top-20 ranking combined with Mavg–Mzmp scatter patterns.

### 2.5. Software implementation and computational performance

MMMAS v1.0.0 is written in Python with a modular architecture. It auto-detects SSR and SNP genotype formats and common missing-value encodings without manual configuration; SNP analysis currently supports format detection and matrix construction, while rate-based thresholds for high-density panels remain under development (Section 4.5). The computation engine selects among nested-loop, NumPy-vectorized, and chunked multi-threaded strategies according to sample size. Before downstream analysis, every run applies five mathematical validation checks to the mismatch matrix (symmetry, non-negativity, boundedness, integrality, and the diagonal rule). Two interfaces are provided: a bilingual (Chinese/English) graphical interface for germplasm curators and a command-line interface for batch processing and pipeline integration. The input is a standard comma-separated values (CSV) genotype matrix; each complete run yields the core matrices, MES grading, duplicate detection, and summary reports. The software is openly available under the MIT license at https://github.com/MMMsystem/MMMAS, with the archived version on Zenodo (https://doi.org/10.5281/zenodo.21441606).

Benchmarks were run on two workstations (Table 1): an ordinary workstation (W1: Windows 10, Intel quad-core 3.3 GHz, 12 GB RAM) and a high-performance workstation (W2: Windows 11, AMD 16-core 2.5 GHz, 32 GB RAM). The simulated benchmark dataset (10,000 accessions × 30 SSR loci) was generated in R using a custom script (available in the MMMAS repository): allele pools comprised 11–26 alleles per locus with gamma-distributed frequencies (shape = 1.2) sampled under Hardy–Weinberg proportions, combining 8,000 unrelated base accessions (5,500 cultivated accessions from five geographic gene pools, 1,500 wild and 1,000 related-population accessions) and 2,000 accessions with embedded known relationships (parent–offspring trios and pairs, grandparent–grandchild, full-sib, half-sib, open-pollinated and distant-cross progeny), with a per-transmission mutation rate of 0.001 and 3% missing genotypes (3.6% realized). All runs completed in seconds to minutes even on the ordinary workstation (Table 1). Runtime scaled nearly linearly with the number of pairs: an 85-fold increase in pairs on W1 produced an 89-fold increase in runtime, as expected for O(N²L) pairwise evaluation. Matrix construction accounted for only about 1% of each run; most runtime was spent on MES grading and report generation.

**Table 1.**
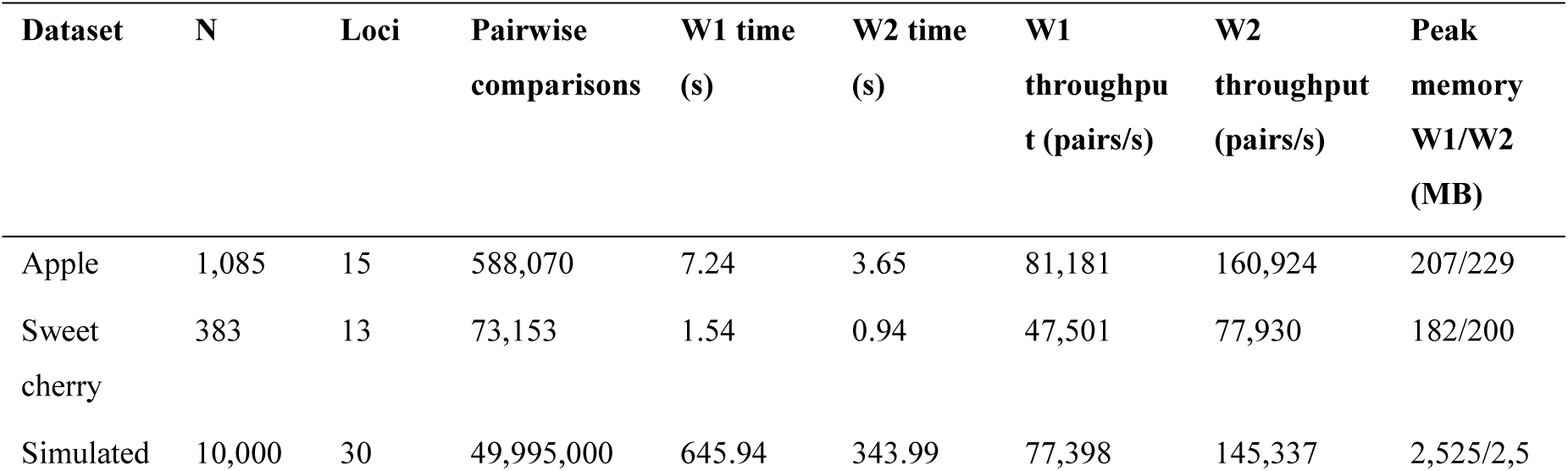
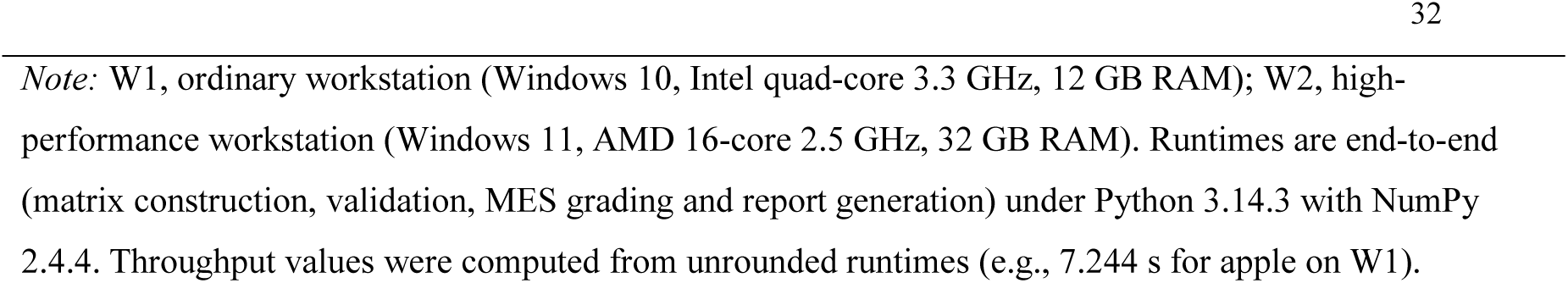
Performance benchmarks of MMMAS v1.0.0 on two workstations.

### 2.6. Data analyses

Descriptive statistics were computed from the MMMAS output matrices using Python (NumPy); figures were prepared with Origin 2024.

## 3. Results

### 3.1. Core index distributions

The mismatch matrices comprised 588,070 independent pairwise comparisons for apple and 73,153 for sweet cherry (Table 2). The two species showed contrasting patterns (Fig. 1). Apple had an approximately symmetric Mmn distribution: only approximately 0.20% of pairs showed zero mismatches (n = 1,200) and approximately 0.47% showed ≤ 1 (n = 2,782). Sweet cherry was right-skewed: 3.12% of pairs had zero mismatches (n = 2,283) and 13.27% had ≤ 1 (n = 9,710). Individual-level indices mirrored this contrast: Mmin values were concentrated at zero (73.73% of apple and 89.82% of sweet cherry accessions), and Mzmp was strongly right-skewed in both species, with higher overall connectivity in sweet cherry (mean 11.92 vs. 2.21) (Fig. 1c–h). The core indices thus revealed two population structures with distinct management implications: a highly diverse, gradient-type structure in apple and a low-diversity, highly homogenized structure in sweet cherry, the latter consistent with the founder-clone bottleneck documented for modern cultivars (Mariette *et al*. 2010).

**Fig. 1.**
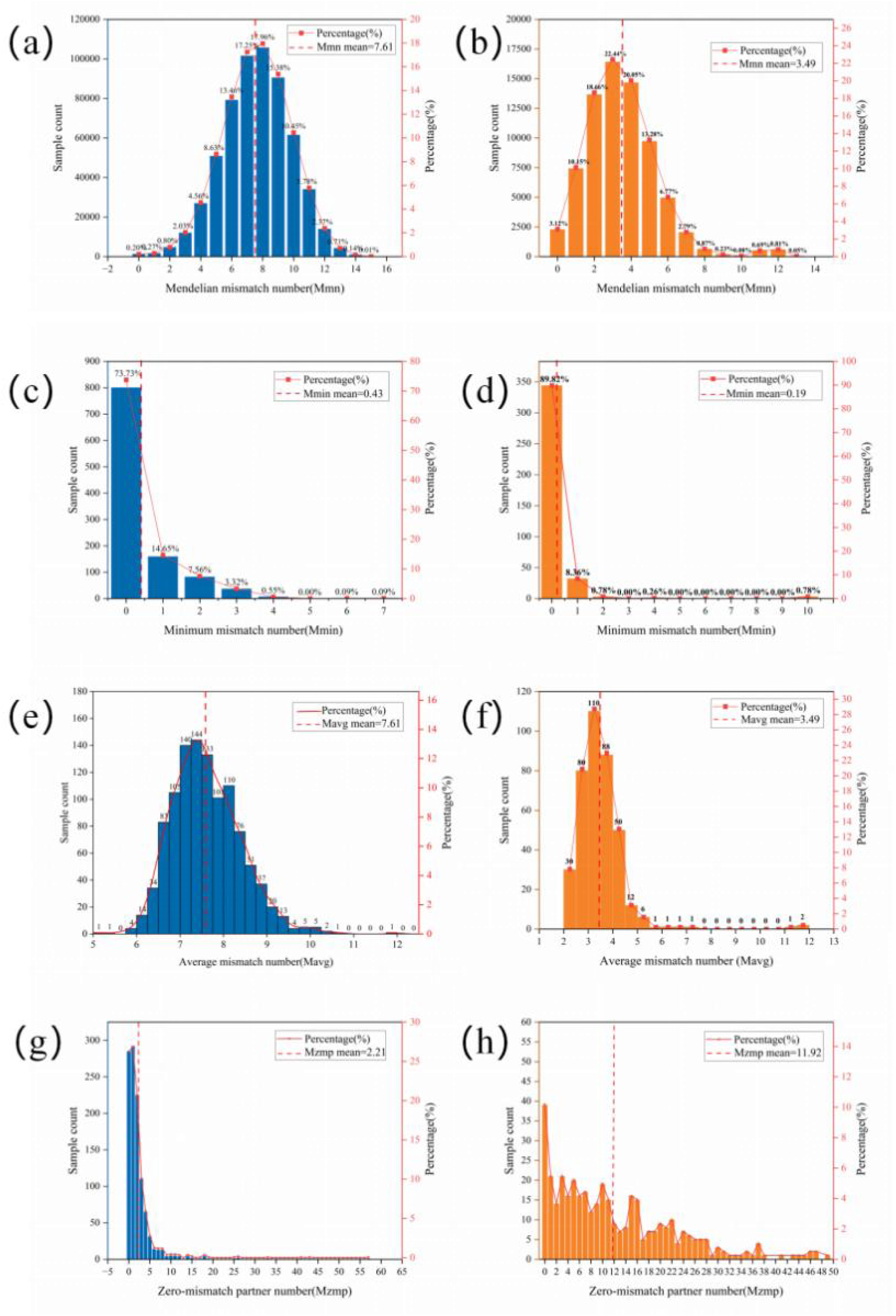
Frequency distributions of four core MMM indices. (a, c, e, g) Apple (N = 1,085, 15 SSRs); (b, d, f, h) sweet cherry (N = 383, 13 SSRs). (a, b) Population-level Mendelian mismatch number (Mmn); (c, d) Mendelian minimum mismatch number (Mmin); (e, f) Mendelian average mismatch number (Mavg); (g, h) Mendelian zero-mismatch partner number (Mzmp).

**Table 2.**
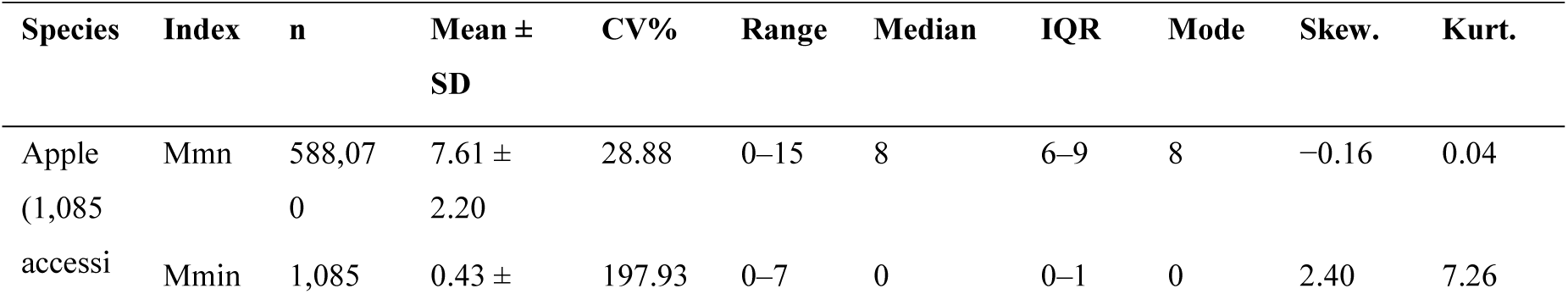
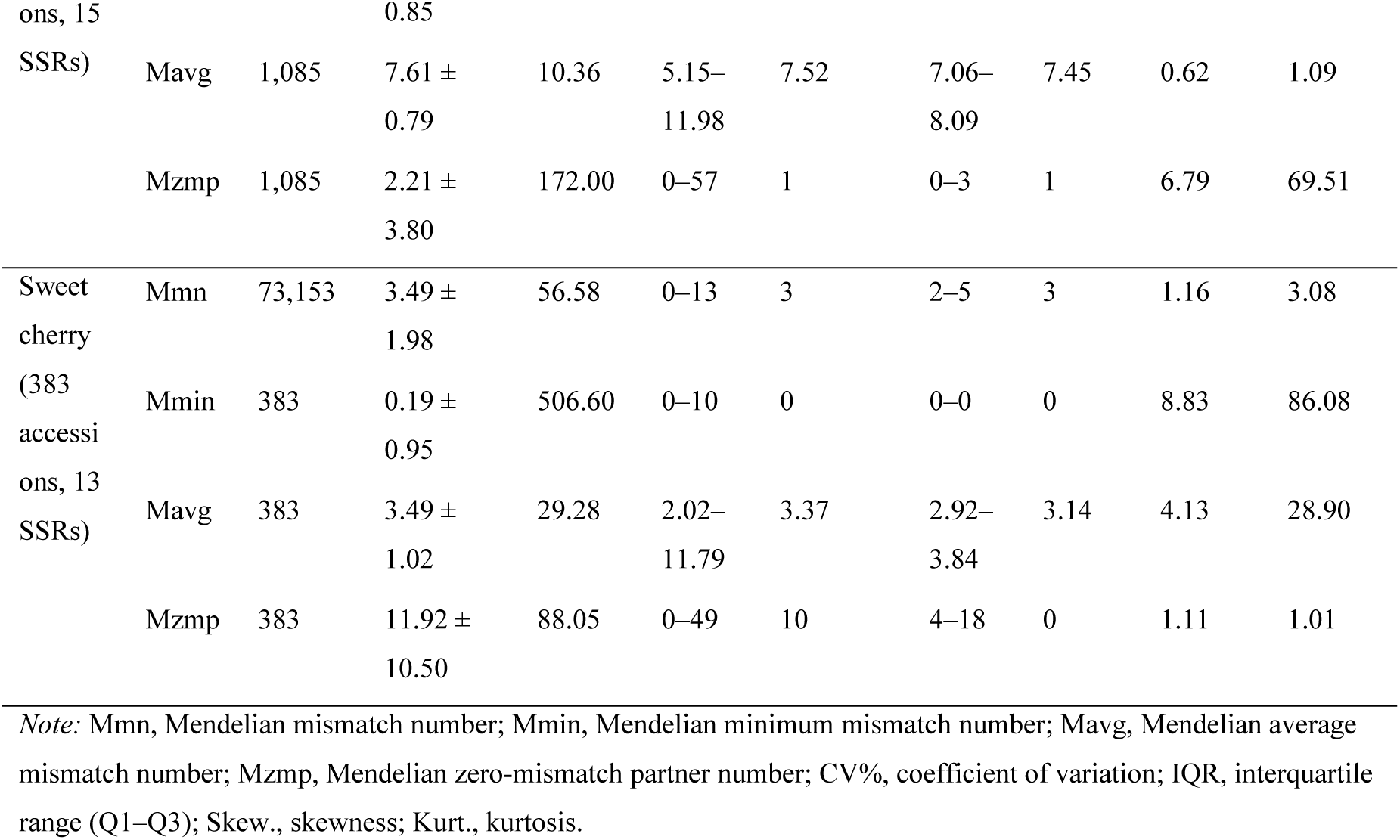
Statistical description of core MMM indices for apple and sweet cherry.

### 3.2. MES stratification

MES stratification quantified these structures (Table 3). Apple displayed a gradient-type hierarchy (MGI = 0.125): grade D₀ contained 800 accessions (73.73%), grade sizes contracted smoothly, and grades D₄–D₇ jointly comprised only eight accessions (0.74%); the wild crabapple *Malus ×robusta* 5 was assigned to D₆. Sweet cherry exhibited a broken-type hierarchy (MGI = 0.545): D₀ contained 344 accessions (89.82%), grades D₃ and D₅–D₉ were all empty, and D₁₀ abruptly contained three accessions (0.78%), all wild species (*Prunus incisa* E621, *P. mahaleb* SL 64 and *P. nipponica* F1292). Such fragmentation is expected given the extreme homogenization of cultivated sweet cherry after a founder-clone bottleneck (Mariette *et al*. 2010).

**Table 3.** MES grade distributions in apple and sweet cherry.

| Species | Grade<br>(Mmin =<br>k) | C-grade<br>n (≥ k) | C-grade % | D-grade<br>n (= k) | D-grade % | Cumulati<br>ve % | Mavg<br>Mean ±<br>SD | Mavg<br>Range |
| --- | --- | --- | --- | --- | --- | --- | --- | --- |
| Apple | 0 | 1,085 | 100.00 | 800 | 73.73 | 73.73 | 7.52 ±<br>0.74 | 5.15–<br>10.68 |
|  | 1 | 285 | 26.27 | 159 | 14.65 | 88.39 | 7.56 ±<br>0.77 | 6.03–<br>10.00 |
|  | 2 | 126 | 11.61 | 82 | 7.56 | 95.94 | 8.03 ±<br>0.73 | 6.26–<br>10.47 |
|  | 3 | 44 | 4.06 | 36 | 3.32 | 99.26 | 8.52 ±<br>0.48 | 7.55–9.30 |
|  | 4 | 8 | 0.74 | 6 | 0.55 | 99.82 | 9.03 ±<br>0.34 | 8.72–9.62 |
|  | 5 | 2 | 0.18 | 0 | 0.00 | 99.82 | – | – |
|  | 6 | 2 | 0.18 | 1 | 0.09 | 99.91 | 10.06 | 10.06 |
|  | 7 | 1 | 0.09 | 1 | 0.09 | 100.00 | 11.98 | 11.98 |
| Sweet<br>cherry | 0 | 383 | 100.00 | 344 | 89.82 | 89.82 | 3.31 ±<br>0.60 | 2.02–5.41 |
|  | 1 | 39 | 10.18 | 32 | 8.36 | 98.17 | 4.41 ±<br>0.58 | 3.62–6.11 |
|  | 2 | 7 | 1.83 | 3 | 0.78 | 98.96 | 5.66 ±<br>0.77 | 5.18–6.55 |
|  | 3 | 4 | 1.04 | 0 | 0.00 | 98.96 | – | – |

| Species | Grade<br>(Mmin =<br>k) | C-grade<br>n (≥ k) | C-grade % | D-grade<br>n (= k) | D-grade % | Cumulative % | Mavg<br>Mean ±<br>SD | Mavg<br>Range |
| --- | --- | --- | --- | --- | --- | --- | --- | --- |
|  | 4 | 4 | 1.04 | 1 | 0.26 | 99.22 | 7.46 | 7.46 |
|  | 5–9 | 3 | 0.78 | 0 | 0.00 | 99.22 | – | – |
| | 10 | 3 | 0.78 | 3 | 0.78 | 100.00 | 11.57 ±<br>0.25 | 11.29–<br>11.79 |

Note: C(k), cumulative grade (accessions with Mmin ≥ k); D(k), difference grade (accessions with Mmin = k). Apple MGI = 0.125; sweet cherry MGI = 0.545. Sweet cherry grades D<sub>5</sub>–D<sub>9</sub> are collapsed for compactness (all empty, C n = 3).

### 3.3. Genetically distinct accessions

Path A (Mmin ≥ 4) identified eight distinct apple accessions and four distinct sweet cherry accessions (Table 4). The apple set included two wild relatives contributing known resistance genes: *Malus ×floribunda* 821 at D₇ (donor of the Rvi6 scab resistance gene; Bus *et al*. 2011) and *M. ×robusta* 5 at D₆ (fire blight resistance; Peil *et al*. 2007; Fahrentrapp *et al*. 2013); the three sweet cherry D₁₀ accessions were all wild relatives. Path B (Tukey outliers on Mavg) complemented the local criterion with a global measure of distinctness: the apple threshold of 9.64 flagged 14 accessions (1.29%), of which 12 (85.71%) fell in grades D₀–D₂, and the sweet cherry threshold of 5.22 flagged 10 accessions, of which six fell in D₀–D₂ (Fig. 2), confirming that low local distinctness does not imply low global distinctness. After de-duplication, the two paths jointly covered 20 apple and 10 sweet cherry highly distinct accessions.

**Fig. 2.**
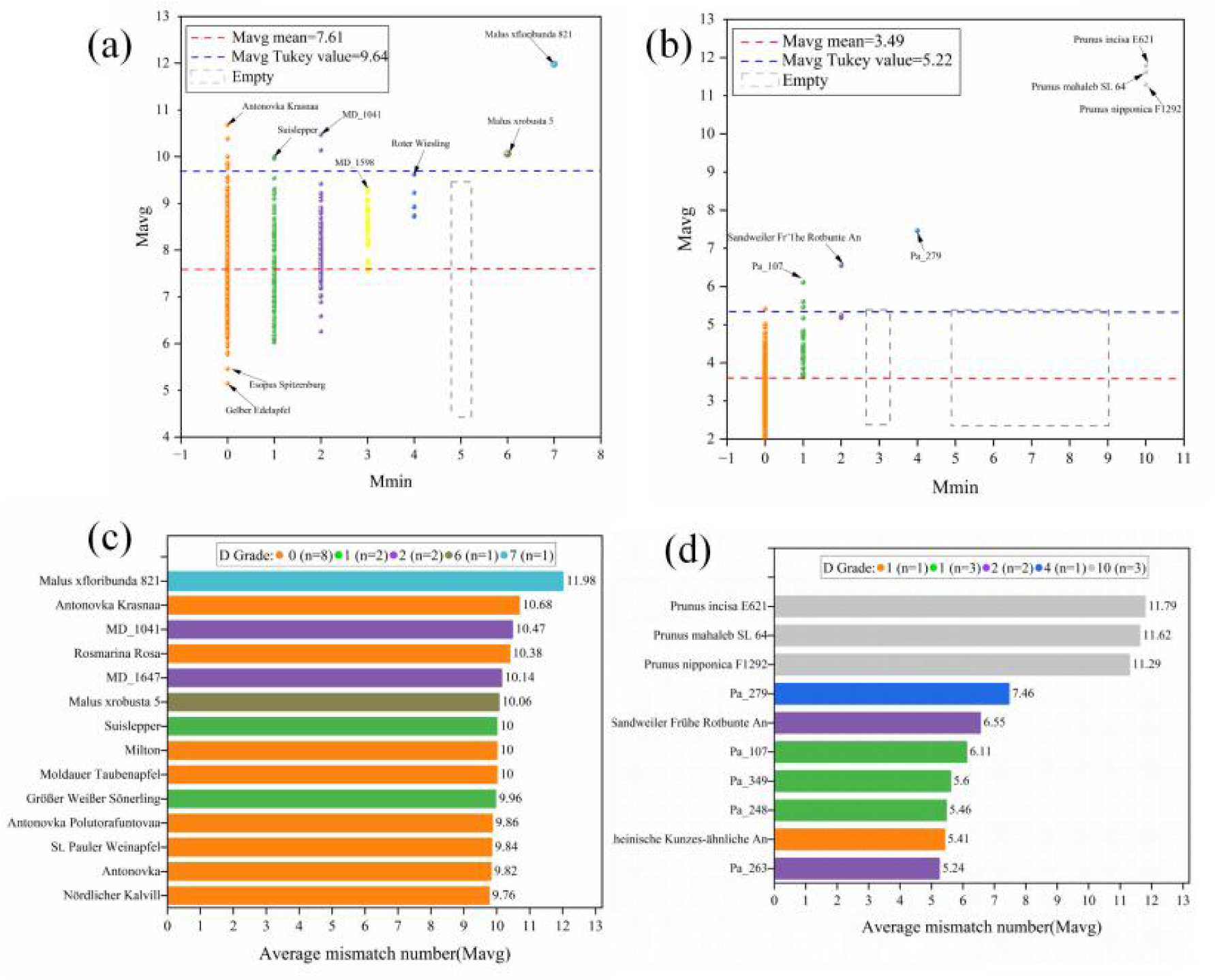
Mmin–Mavg dual-index joint strategy for identifying genetically distinct germplasm. (a) Apple dual-index distribution with the Tukey threshold (9.64) as a blue dashed line; (b) 14 highly globally distinct apple accessions; (c) sweet cherry dual-index distribution with the Tukey threshold (5.22); (d) highly globally distinct sweet cherry accessions.

**Table 4.** High-grade D₄⁺ distinct accessions in apple and sweet cherry.

| Species | MES grade | Accession | Mavg |
| --- | --- | --- | --- |
| Apple | D7 | *Malus* × *floribunda* 821 | 11.98 |
| Apple | D6 | *Malus* × *robusta* 5 | 10.06 |
| Apple | D4 | Roter Wiesling | 9.62 |
| Apple | D4 | Trübchen An | 9.23 |
| Apple | D4 | Himbsels Rambur | 8.93 |
| Apple | D4 | Batullenapfel | 8.92 |
| Apple | D4 | Engelshofer | 8.74 |
| Apple | D4 | Zelenka | 8.72 |
| Sweet cherry | D10 | *Prunus incisa* E621 | 11.79 |
| Sweet cherry | D10 | *P. mahaleb* SL 64 | 11.62 |
| Sweet cherry | D10 | *P. nipponica* F1292 | 11.29 |
| Sweet cherry | D4 | Pa_279 | 7.46 |

### 3.4. Genetic hub accessions

Mzmp rankings closely matched the documented breeding history of both species (Table 5, Fig. 3). In apple, top-ranked accessions were led by ‘Cox Orange’ (Mzmp = 57). High-Mzmp individuals included founding clones of modern apple cultivars identified by Noiton and Alspach (1996), such as ‘Cox Orange’, ‘Golden Delicious’, ‘Jonathan’, ‘McIntosh’, and ‘Delicious’, together with breeding-hub parents subsequently verified through molecular markers or pedigree reconstruction, such as ‘James Grieve’ (Larsen *et al*. 2024; Muranty *et al*. 2020; Howard *et al*. 2021b; Luby *et al*. 2022). This outcome is consistent with large-scale studies of founding clones and pedigree structure in apple germplasm (Urrestarazu *et al*. 2016; Vanderzande *et al*. 2019; Muranty *et al*. 2020), indicating that Mzmp identifies centers of genetic connectivity.

**Fig. 3.**
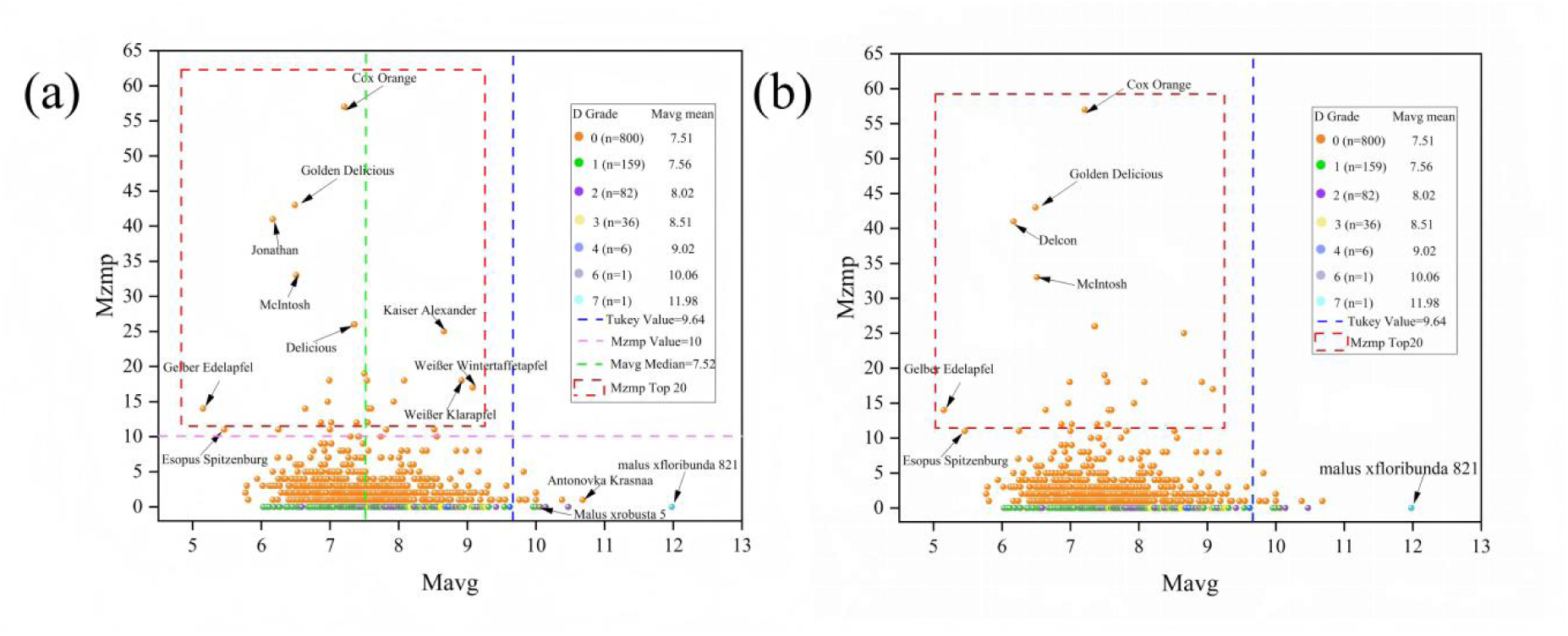
Mavg–Mzmp dual-index scatter plots for apple (a) and sweet cherry (b). Blue dashed lines denote the Tukey thresholds (9.64 for apple; 5.22 for sweet cherry); green dashed lines denote the Mavg median for apple (7.52); pink dashed lines denote the Mzmp = 10 threshold for apple; red dashed boxes highlight the top 20 hubs ranked by Mzmp.

**Table 5.** Top-ranking hub accessions by Mzmp.

| Rank | Apple | Mzmp | Mavg | Sweet cherry | Mzmp | Mavg |
| --- | --- | --- | --- | --- | --- | --- |
| 1 | **Cox Orange** | 57 | 7.21 | **Lübecker Bunte** | 49 | 2.32 |
| 2 | **Golden Delicious** | 43 | 6.49 | **Dudenroter Knorpelkirsche** | 47 | 2.12 |
| 3 | **Jonathan** | 41 | 6.17 | **Pa_223** | 47 | 2.50 |
| 4 | **McIntosh** | 33 | 6.51 | **Bopparder Krächer** | 46 | 2.18 |
| 5 | **Goldparmäne** | 26 | 7.35 | **Reiffenhäuser Herzförmige An** | 46 | 2.26 |
| 6 | **Delicious** | 26 | 7.36 | **Große Schwarze Knorpelkirsche** | 45 | 2.02 |
| 7 | **Kaiser Alexander** | 25 | 8.66 | **Knauffs Schwarze** | 44 | 2.37 |
| 8 | **James Grieve** | 19 | 7.50 | **Badacsoner aus Adocs** | 43 | 2.10 |
| 9 | **Boikenapfel** | 18 | 8.08 | **Kunzes Kirsche** | 41 | 2.07 |
| 10 | Kasseler Renette | 18 | 7.54 | Volltragende Knorpelkirsche | 38 | 2.36 |
| 11 | Prinzenapfel | 18 | 6.99 | Elkershäuser Schwarzkirsche An | 37 | 2.32 |
| 12 | Weißer Klarapfel | 18 | 8.92 | Hedelfinger Riesenkirsche | 37 | 2.53 |
| 13 | Weißer Wintertaffetapfel | 17 | 9.08 | Späte Rundliche | 37 | 2.19 |
| 14 | Bismarckapfel | 15 | 7.93 | Späte Spanische | 37 | 3.67 |
| 15 | Orleans Renette | 15 | 6.97 | Ria | 36 | 2.02 |
| 16 | Gelber Edelapfel | 14 | 5.15 | Pa_053 | 35 | 2.49 |
| 17 | Sunset | 14 | 7.56 | Spanische vom Mittelrhein | 35 | 2.84 |
| 18 | Clivia | 14 | 6.64 | Beata | 34 | 2.40 |
| 19 | Wealthy | 14 | 7.60 | Große Prinzessin | 33 | 2.77 |
| 20 | Weißer Winterkalvill | 12 | 7.55 | Techlovan | 32 | 2.66 |
| 20† | Geheimrat Dr. Oldenburg | 12 | 7.02 |  |  |  |
| 20† | Worcester Parmène | 12 | 7.39 |  |  |  |
| 20† | Danziger Kantapfel | 12 | 6.87 |  |  |  |

Note: † indicates tied at rank 20 (Mzmp = 12). For sweet cherry, the rank grouping follows documented breeding history: ranks 1 – 10 are ancient founder cultivars and landraces, and ranks 11 – 20 are modern breeding parents with marker-verified pedigrees.

The sweet cherry top 20 showed a two-tier structure (Table 5). Ranks 1–10 (Mzmp = 38–49), led by ‘Lübecker Bunte’ (49), were ancient founder cultivars and landraces without breeding records, such as ‘Bopparder Krächer’ (46). Ranks 11–20 (Mzmp = 32–37) comprised modern breeding parents with marker-supported parentage, such as ‘Hedelfinger Riesenkirsche’ and ‘Techlovan’ (Quero-García *et al*. 2017; Reim *et al*. 2023; Choi and Kappel 2004).

Mzmp also uncovered structurally central accessions absent from the pedigree literature. The unnamed sweet cherry accession Pa_223 (Mzmp = 47, jointly the second highest and behind only ‘Lübecker Bunte’ (49)) had zero-mismatch partners spanning ancient landraces and modern cultivars of diverse geographic origins. In apple, high-Mzmp accessions without documented breeding records included ‘Goldparmäne’ (Mzmp = 26) and ‘Kaiser Alexander’ (Mzmp = 25) (Table 5). International molecular surveys have revealed previously unknown heteroploid and inbred relationships (Ordidge *et al*. 2018), so many genetic connections plausibly remain undocumented. Highly ranked accessions without published records may therefore represent unrecognized parental hubs or diversity reservoirs; unlike pedigree analysis, Mzmp requires no pedigree information.

### 3.5. Functional stratification of germplasm

Accessions were classified into four management-relevant functional quadrants from the joint Mavg–Mzmp distribution of the apple collection, using the dataset-specific thresholds defined in Section 2.2: Mzmp ≥ 10 (capturing the top 3.0% of connectivity), the Mavg median (7.52), and the Tukey upper fence (9.64) (Table 6; Fig. 3a). Bridge accessions (n = 13) combine high connectivity with high average genetic distance; the most extreme example, ‘Kaiser Alexander’ (Mzmp = 25, Mavg = 8.66), has partners spanning distinct European apple traditions, and such accessions are candidate bridging parents for wide crosses and introgression breeding. Pedigree cores (n = 20) are documented founding parents such as ‘Cox Orange’ (57, 7.21) and ‘Golden Delicious’ (43, 6.49), whose partners concentrate in genetically compact neighborhoods. Genetic extremes (n = 14) include the wild relatives *Malus ×floribunda* 821 (0, 11.98) and *M. ×robusta* 5 (0, 10.06) as well as the isolated cultivar ‘Antonovka Krasnaa’ (1, 10.68), meriting the highest conservation priority and whole-genome sequencing. Standard accessions (n = 1,038) form the network background and require periodic redundancy assessment.

**Table 6.**
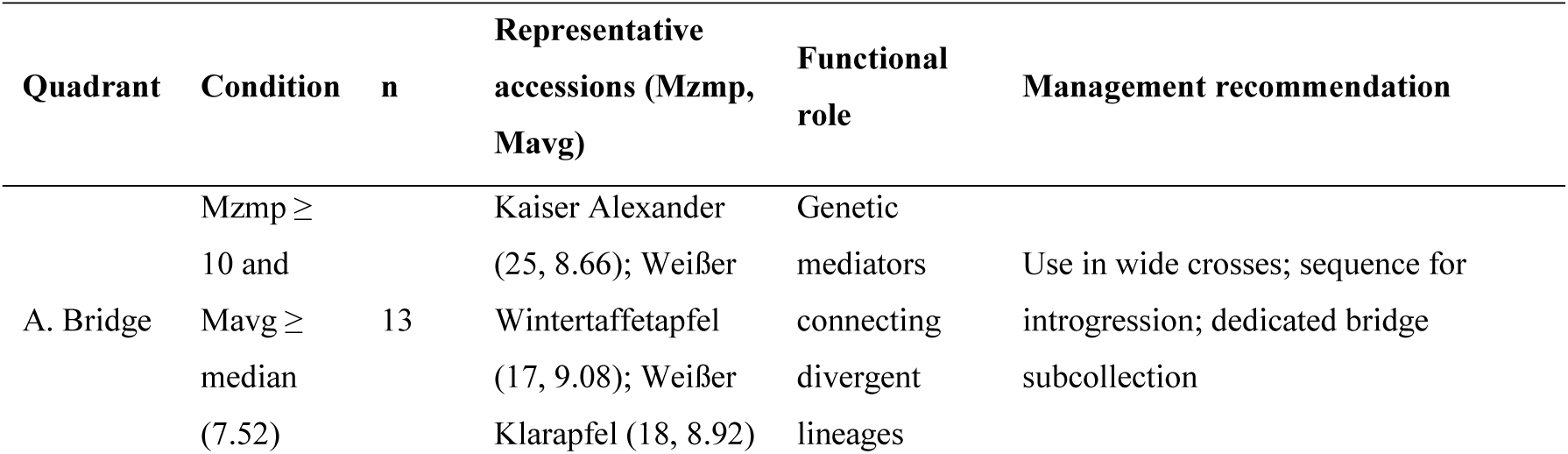
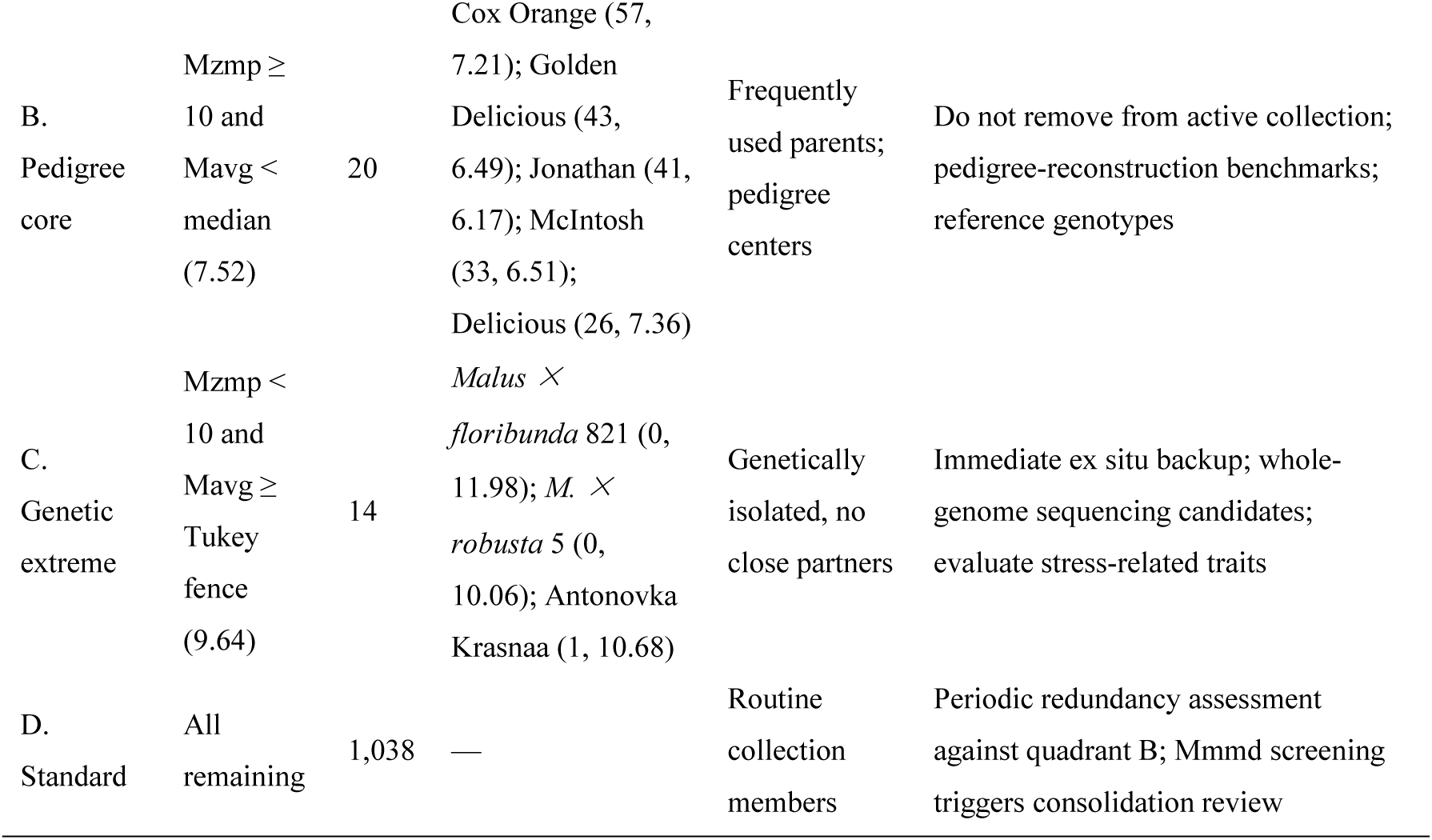
Functional quadrant classification of the apple collection (N = 1,085, de-duplicated), representative accessions, and management recommendations.

The framework was applied to the apple collection, where the broad Mavg–Mzmp contrast provides clear functional discrimination. In the highly homogenized sweet cherry collection (Section 3.1), the narrow Mavg distribution and uniformly high connectivity leave limited contrast for quadrant separation; functional identification there was achieved instead through Mzmp ranking, which uncovered the undocumented hub accession Pa_223. The quadrant classification thus links genotype-based stratification directly to conservation and breeding decisions (Table 6).

### 3.6. Pedigree concordance

Among the 335 CERVUS-confirmed apple parent–offspring pairs, 97.31% (326/335) fell within Mmn ≤ 1 (82.09% with Mmn = 0; 15.22% with Mmn = 1), and only 2.69% (9/335) had Mmn ≥ 2, on the same order as reported SSR genotyping error rates (Bonin *et al*. 2004; Wang 2010). Conversely, 97.83% (45/46) of CERVUS-rejected pairs had Mmn ≥ 2; mean mismatch counts were 0.24 for confirmed versus 6.30 for rejected pairs (Table 7). Precision among the 327 CERVUS-evaluated Mmn ≤ 1 candidates was 99.69% (326/327); the wider MMM candidate set (2,782 pairs) reflects the intended behavior of a coarse screen. In sweet cherry, 75.86% (66/87) of evaluable pairs had Mmn = 0 and 22.99% (20/87) had Mmn = 1; after stratification by confidence, all 83 pairs with confidence ≥ 80% fell within Mmn ≤ 1 (100% concordance). The single high-mismatch case (Nafrina × Büttners Rote Knorpelkirsche, Mmn = 4) was also the only CERVUS-evaluated trio with confidence below 80%; these two lines of evidence corroborate each other (Reim *et al*. 2023). Among the remaining low-confidence records, the ‘Katalin’ mother–offspring pair (Mmn = 0) and the ‘Swing’ father–offspring pair (Mmn = 1) still fell within the low-mismatch range.

**Table 7.**
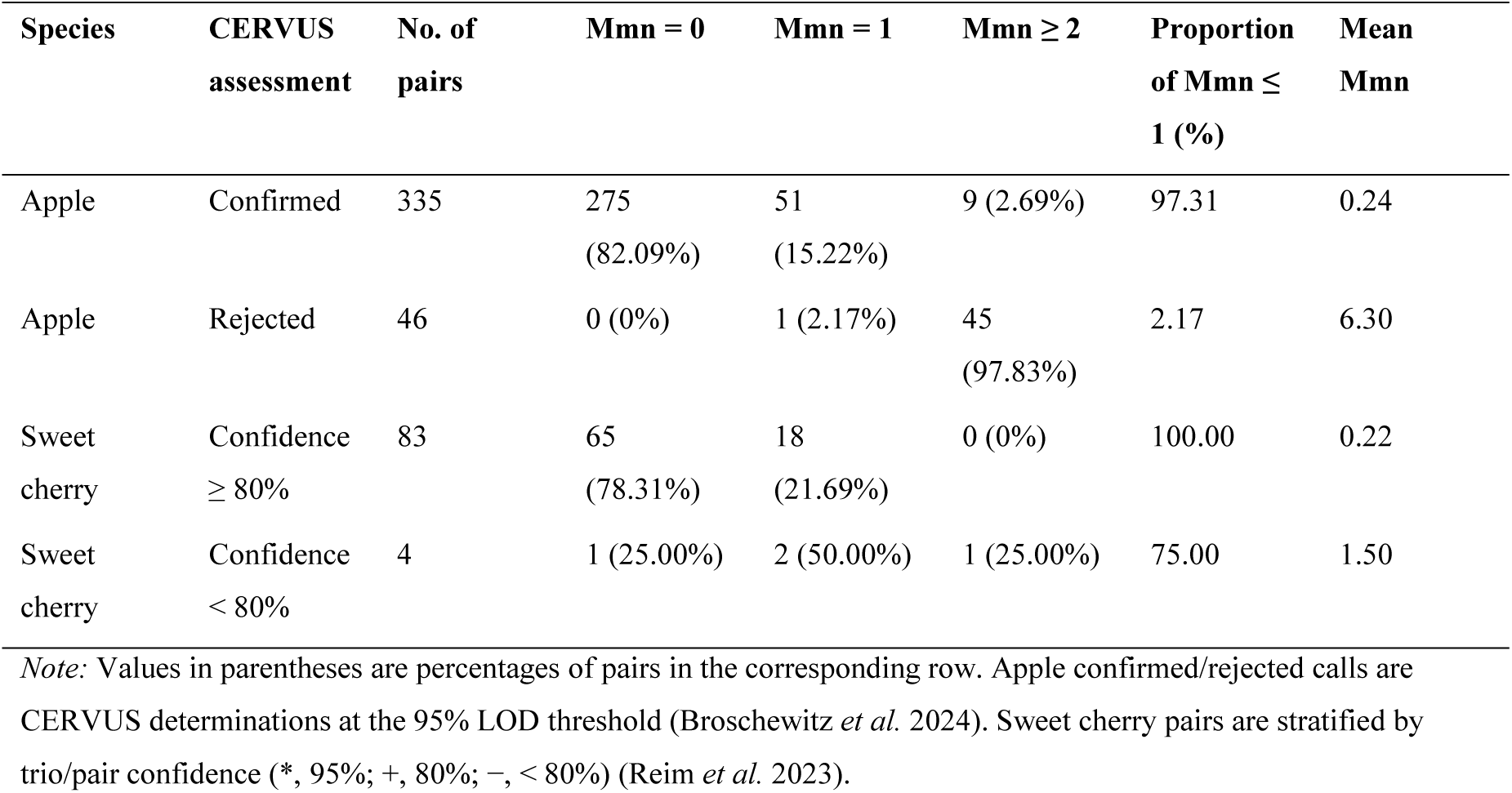
Concordance between Mmn and published CERVUS pedigree assessments.

| Species | CERVUS<br>assessment | No. of<br>pairs | Mmn = 0 | Mmn = 1 | Mmn ≥ 2 | Proportion<br>of Mmn ≤<br>1 (%) | Mean<br>Mmn |
| --- | --- | --- | --- | --- | --- | --- | --- |
| Apple | Confirmed | 335 | 275<br>(82.09%) | 51<br>(15.22%) | 9 (2.69%) | 97.31 | 0.24 |
| Apple | Rejected | 46 | 0 (0%) | 1 (2.17%) | 45<br>(97.83%) | 2.17 | 6.30 |
| Sweet<br>cherry | Confidence<br>≥ 80% | 83 | 65<br>(78.31%) | 18<br>(21.69%) | 0 (0%) | 100.00 | 0.22 |
| Sweet<br>cherry | Confidence<br>< 80% | 4 | 1 (25.00%) | 2 (50.00%) | 1 (25.00%) | 75.00 | 1.50 |

Note: Values in parentheses are percentages of pairs in the corresponding row. Apple confirmed/rejected calls are CERVUS determinations at the 95% LOD threshold (Broschewitz *et al.* 2024). Sweet cherry pairs are stratified by trio/pair confidence (\*, 95%; +, 80%; -, < 80%) (Reim *et al.* 2023).

### 3.7. Duplicate detection

Among the 1,200 zero-mismatch candidate pairs in the apple dataset, no Mmmd = 1 pair was detected, consistent with the prior genotype de-duplication of that collection (Broschewitz *et al*. 2024). In sweet cherry, exactly one Mmmd = 1 pair was detected (‘Büttners Rote Knorpelkirsche’ and ‘Querfurter Königskirsche’), a pair confirmed as duplicate genotypes in the literature (Reim *et al*. 2023). The Mmmd index is computationally simple, yields unambiguous decisions, and is well suited as a rapid duplicate-checking step at accession intake.

### 3.8. Marker-panel sensitivity

Reducing the apple marker panel produced three distinct patterns (Fig. 4). Gradient compression: the proportion of D₀ accessions rose from 63.04% (17 SSRs) to 97.97% (5 SSRs), with two accelerating phases: the null-allele filtering step (17→15, +10.69 pp) and the low-density end (9→7, +8.11 pp, 85.44%→93.55%). Grade truncation: the highest attainable grade fell from D₉ to D₁, and D₄⁺ accessions dropped from 47 (17 SSRs) to 8 (15 SSRs, −83.0%) and 2 (13 SSRs), reaching zero at 9 SSRs. The two wild, disease-resistant accessions illustrate the cost of low-density panels: *M. ×floribunda* 821 fell from D₉ (17 SSRs) to D₀ (5 SSRs), a complete loss of distinctness recognition, entering the D₀ class with three zero-mismatch partners at 5 SSRs; *M. ×robusta* 5 declined from D₇ (17 SSRs) to D₁ (5 SSRs). Collections conserving wild relatives or resistance donors should therefore use no fewer than 13 loci; on this dataset, 11 high-PIC loci (mean PIC = 0.837) sufficed for stable D₄⁺ identification, the two D₄⁺ accessions at 13 loci (*M. × robusta* 5 and *M. ×floribunda* 821) being the same two at 11 loci. The Mzmp of ‘Cox Orange’ rose monotonically as the panel narrowed (45 at 17 SSRs; 57 at 15, 13 and 11; 61 at 9; 65 at 7; 68 at 5). Low-resolution panels therefore overestimate the Mendelian compatibility radius of core parents (68 at 5 SSRs versus 45 at 17 SSRs). Mmmd stability: the index requires sufficient marker density to maintain vector resolution. In this pre-de-duplicated collection, 17→9 SSR panels yielded no duplicate groups, whereas reducing to 7 SSR flagged 1 spurious pair and reducing to 5 SSR flagged 2 spurious pairs, reflecting loss of vector resolution rather than genuine duplication.

**Fig. 4.**
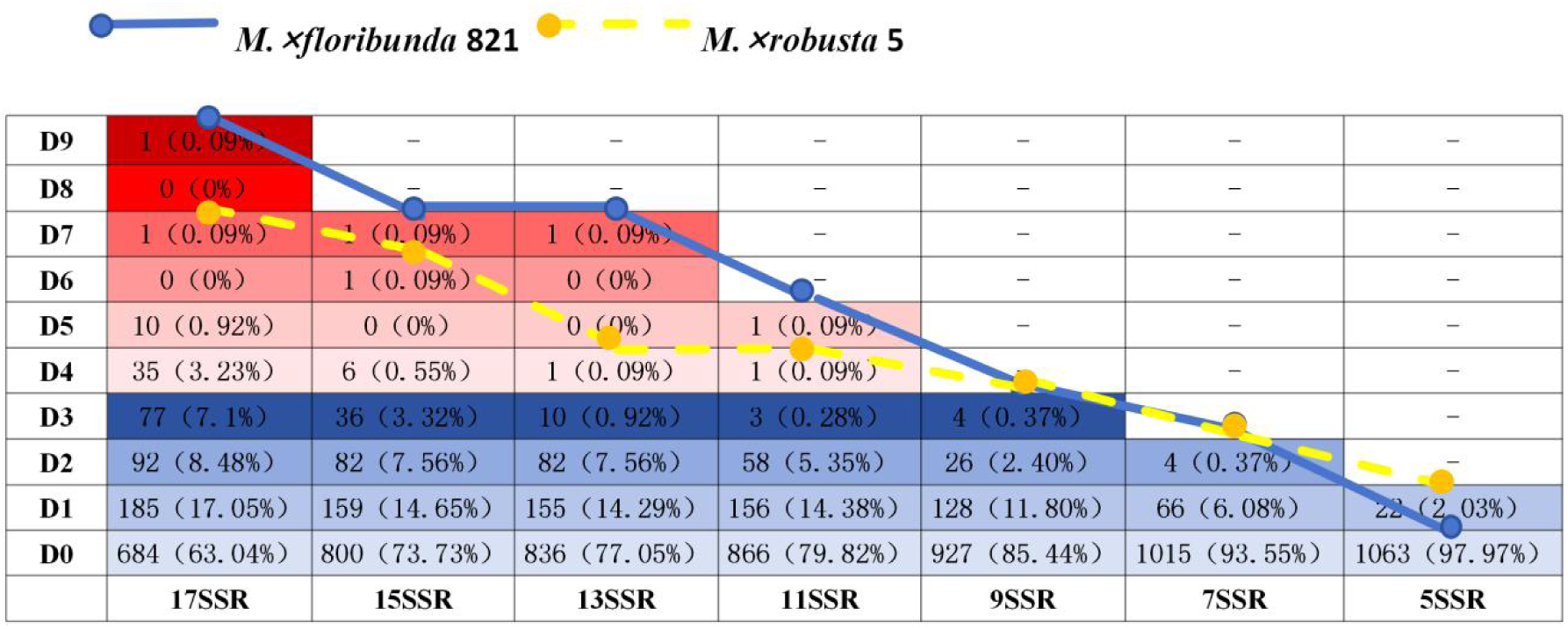
Distribution of MES difference grades (D₀–D₉) under progressive SSR locus reduction from 17 to 5. Cell values indicate accession counts with proportions in parentheses; warm colors denote D₄–D₉, cool colors denote D₀–D₃. The blue solid line traces *Malus ×floribunda* 821 and the yellow dashed line traces *M. ×robusta* 5, positioned at D₉ and D₇ with 17 SSRs; both accessions shift toward lower grades as marker density decreases, with *M. ×floribunda* 821 falling to D₀ (complete loss of distinctness recognition) and *M. ×robusta* 5 to D₁ at 5 SSRs.

## 4. Discussion

### 4.1. Positioning MMMAS in the germplasm management toolkit

The MMM framework complements existing horticultural genomics tools at multiple levels. While pan-transcriptome studies in pear have captured transcriptome-level diversity (Sun *et al*. 2025) and high-throughput SNP arrays such as Melon2K provide cost-effective genotyping platforms for molecular breeding (Yu *et al*. 2025b), these tools focus primarily on diversity discovery rather than on downstream quality management. MMM fills this gap by converting raw genotype data into curation decisions for collections of thousands of accessions.

MMM is implemented as the open-source software MMMAS, a deterministic pre-screening framework that requires neither allele-frequency estimates nor genetic model assumptions. Unlike distance-based clustering methods that depend on arbitrary cutoff parameters, MMM builds on the discrete nature of Mendelian exclusion: each SSR locus either excludes a parent–offspring relationship or does not, so the resulting mismatch grades reflect biological reality directly. This deterministic logic shares conceptual roots with the opposing-homozygote approach used for pedigree verification in livestock (Calus *et al*. 2011) but generalizes it to arbitrary pairwise relationships and population-level analyses without requiring known pedigree structures.

The framework derives collection-scale diagnostics (Mmin, Mavg, Mzmp, Mmmd, and MES) from a single matrix in one pass, rather than returning pairwise relationship assessments, addressing a known limitation of germplasm management workflows. Existing parentage tools such as CERVUS (Kalinowski *et al*. 2007), hiphop (Cockburn *et al*. 2021), COLONY (Jones and Wang 2010) and Sequoia (Huisman 2017) either score candidate parent–offspring pairs or cluster sibships and pedigrees, yet each returns relationship-level assignments rather than a collection-wide synthesis. Genome-wide relatedness tools such as PLINK (Purcell *et al*. 2007), KING (Manichaikul *et al*. 2010), and GCTA (Yang *et al*. 2011) compute genome-wide similarity matrices that encode continuous allele-sharing coefficients rather than discrete Mendelian exclusions, reflecting the broader shift of parentage and kinship analysis toward high-density markers (Flanagan and Jones 2019). Core-collection tools such as PowerCore (Kim *et al*. 2007) and Core Hunter (De Beukelaer *et al*. 2018) maximize allelic diversity with minimal redundancy, but report neither duplicate genotype pairs nor mislabeled accessions.

In contrast to SNP-based pedigree reconstruction, which requires genome-wide data and empirically derived error thresholds (Muranty *et al*. 2020; Yang *et al*. 2022), MMM operates on 10–20 SSR loci using exact Mendelian exclusion. Howard *et al*. (2021b) showed that shared ploidy-adjusted haplotype length (SPLoSH) information enables pedigree reconstruction in outbreeding crops. Although SPLoSH requires genome-wide SNP data for haplotype phasing, its outputs (confirmed parent–offspring pairs and trios) represent ideal downstream validation targets for MMM pre-screening. The deterministic MMM layer identifies candidate pairs in seconds; these candidates can then be prioritized for SPLoSH-based deep reconstruction.

Positioned this way, MMM acts as a stable, model-free first filter rather than a replacement for likelihood-based inference. By flagging genetic conflicts, resolving redundant accessions, and delineating patterns of distinctness in a single pass, it narrows the candidate set that downstream likelihood or haplotype-based frameworks then examine in depth (McMaster *et al*. 2025).

### 4.2. Biological meaning of MES stratification

The contrast in MES structure between apple (a continuous gradient, MGI = 0.125) and sweet cherry (a fragmented hierarchy, MGI = 0.545) reflects their distinct evolutionary and breeding histories. The uninterrupted apple hierarchy is consistent with repeated historical gene flow from wild relatives (Cornille *et al*. 2012), whereas the discrete jumps in sweet cherry mirror the founder-clone bottleneck documented in modern cultivars (Mariette *et al*. 2010; Campoy *et al*. 2016). This species-specific signature suggests that MGI could serve as a candidate metric for summarizing how founder effects and breeding history shape collection structure in fruit crops, pending validation on additional species.

Unlike conventional distance-based clustering methods that compress multi-dimensional genetic data into a single metric, the MES grade system preserves the biological interpretability of Mendelian exclusion: each grade increment represents exactly one additional locus at which a parent–offspring relationship is excluded. This one-locus-per-grade semantics keeps the grades readable without dataset-specific parameter calibration, although the grade distribution itself remains specific to the collection and marker panel examined (Section 2.2).

The same exclusion matrix also yields Mzmp, a quantitative, data-driven route to identifying genetic hubs without prior pedigree knowledge. In apple, ‘Cox Orange’ (Mzmp = 57) and ‘Golden Delicious’ (Mzmp = 43) correspond to founding clones previously identified by Noiton and Alspach (1996) as central to modern apple breeding. In sweet cherry, the Mzmp ranking follows the documented split between ancient founder cultivars and landraces (Mzmp = 38–49) and modern breeding parents (Mzmp = 32–37) rather than a sharp numerical gap (Table 5), paralleling the founder-clone bottleneck described above. Because zero-mismatch partners can include parent–offspring, full–sib, and clonally related or synonymously named accessions (Section 2.2), Mzmp in such a highly homogenized collection amplifies breeding centrality and clonal-family size simultaneously; annotating top-ranked accessions against the synonym and pedigree records of Reim et al. (2023) is a natural curation follow-up.

### 4.3. Practical thresholds and management applications

Marker-reduction analysis (Section 3.8) indicates that diagnostic resolution degrades progressively with marker reduction (D₄ ⁺ accessions fell from 47 with 17 SSRs to 8 with 15 SSRs and 2 with 13 SSRs, disappearing at 9 SSRs), that 11 high-PIC loci sufficed for stable identification of distinct accessions, and that collections conserving wild relatives or resistance donors should use no fewer than 13 loci. These thresholds must be recalibrated for other collections, and standardized SSR panels will improve portability: a coded reference-allele set enabled cross-laboratory comparability of grape SSR profiles (This *et al*. 2004), while single-tube multiplex assays are reducing genotyping cost and error in sweet cherry (Čmejlová *et al*. 2026).

Earlier SSR-based studies typically targeted smaller collections with diversity-description aims; Ouni *et al*. (2020), for example, characterized 61 Tunisian pear accessions. The present framework addresses a different setting: quality management in collections of one thousand accessions and above, where pairwise-only workflows become impractical.

In routine management, MMM diagnostics correspond to conservation priorities: D₄⁺ accessions are candidates for core collections; Mzmp hubs provide structured genetic backgrounds for association studies; and Mmmd-flagged duplicate groups trigger consolidation review, reducing redundant conservation costs (Odong *et al*. 2013; Campoy *et al*. 2016; Anglin *et al*. 2025). The Mavg–Mzmp quadrant classification further introduces a functional annotation layer, assigning each accession a management-relevant role and complementing rather than replacing existing core-collection tools such as PowerCore (Kim *et al*. 2007) and Core Hunter 3 (De Beukelaer *et al*. 2018).

Bridge accessions such as ‘Kaiser Alexander’ (Mzmp = 25, Mavg = 8.66) exemplify what we term “dark core” germplasm: accessions that are structurally central to the collection yet absent from the pedigree literature. Because the apple dataset contains no duplicate genotypes (Section 3.7), this connectivity cannot be attributed to clonal replication; the combination of high connectivity with high average genetic distance instead points to a genuine bridging role between divergent lineages. These accessions are candidates for wide crosses and pedigree research. Parallel evidence from strawberry genealogy indicates that a small number of founding ancestors contribute disproportionately to the allelic diversity of modern cultivars (Pincot *et al*. 2021), suggesting that structurally central yet pedigree-undocumented accessions may also occur in other clonally propagated perennial fruit crops; their frequency awaits confirmation by comparable matrix-based screening in additional species.

A second central role of MMM is the recognition of genetically distinct accessions. Two complementary routes identify them: high MES grades (Path A, D₄⁺) capture local distinctness, whereas high Mavg values (Path B) capture global distinctness; together they flagged 20 apple and 10 sweet cherry accessions, including wild relatives carrying known resistance genes such as *Malus × floribunda* 821 (Rvi6) and *M. × robusta* 5 (Section 3.3). Such accessions are immediate priorities for conservation and whole-genome sequencing.

For perennial fruit crops with long juvenile phases, early detection of genetic duplicates and pedigree errors through deterministic pre-screening can reduce field maintenance costs and streamline core-collection construction. China’s national fruit germplasm system maintains more than 30,000 accessions (Liang *et al*. 2025); in such systems, allele frequencies are often unknown for rare landraces, and computational resources can be limited at provincial genebanks; MMMAS runs on an ordinary workstation within seconds (Table 1).

### 4.4. Concordance, error sensitivity, and robustness

The agreement between the MMM mismatch criterion and published CERVUS assignments (Section 3.6) constitutes a same-data cross-method concordance test rather than an independent validation, because both analyses draw on the same German Fruit Genebank genotype datasets; it therefore shows that MMM captures the same parent–offspring relationships as likelihood-based inference, not that it generalizes beyond these data. Independent validation awaits publicly available SSR datasets with pedigrees documented independently of the genotypes analyzed here. The 2.69% of confirmed apple pairs with Mmn ≥ 2 (9/335) most likely reflect genotyping anomalies such as null alleles, allelic dropout, or batch effects: per-locus SSR error rates of 0.1%–1% are sufficient to produce occasional double mismatches across a 15-locus panel (Bonin *et al*. 2004; Hoffman and Amos 2005; Wang 2010). Determinism is therefore conditional on genotyping accuracy: the matrix flags every pair that violates Mendelian transmission under the error-free assumption, yet it cannot separate technical errors from biological exceptions without replicate-based validation (Kan-Lingwood *et al*. 2025) or pedigree follow-up (Cheung *et al*. 2014).

That most confirmed parent–offspring pairs fall at Mmn ≤ 1 is consistent with a systematic-error interpretation of single mismatches. Arias *et al*. (2022) classified Mendelian errors into eight classes, separating systematic errors (e.g., null alleles) from random calling errors. Within this taxonomy, a single mismatch in an otherwise compatible pair is more plausibly attributed to a systematic error, which recurs at the same locus across many pairs, than to a random miscall, which arises independently in single genotypes and shows no such locus-level recurrence.

Genotyping errors occur even in high-quality datasets, and the Boolean MMM framework accommodates them through the Mmn ≤ 1 acceptance rule: a single mismatch is treated as compatible with Mendelian inheritance, and such pairs accounted for 15.22% and 22.99% of confirmed parent–offspring relationships in apple and sweet cherry, respectively (Section 3.6). This built-in tolerance is particularly valuable for routine genebank screening, where replicate genotyping of every accession is often impractical.

### 4.5. Limitations and future directions

The core Mmn computation is parameter-free, whereas the quadrant thresholds are dataset-dependent exploratory values (Section 2.2). The framework was demonstrated on two diploid fruit-tree species, with agreement against published parentage assessments limited to same-data cross-method concordance (Section 4.4), and the functional quadrant classification was developed on the apple collection alone, so recalibration is required before transferring MMMAS to other collections. The current implementation is further restricted to diploid codominant markers, and polyploidy requires dedicated models because allele dropout biases exclusion counting (Howard *et al*. 2021a; Howard *et al*. 2023; Broschewitz *et al*. 2026). Finally, Boolean exclusion is sensitive to genotyping error, so analyses should start from quality-controlled data.

We did not benchmark MMM directly against SNP-based likelihood methods, because the two operate at different stages of the analysis pipeline; a further obstacle is the scarcity of publicly available SNP genotype datasets with documented pedigrees suitable for validation. For SNP-scale application, SSR-derived absolute-count thresholds should not be transferred directly to high-density panels. SNP array data harbor abundant Mendelian errors (null alleles, genomic alterations; Arias *et al*. 2022), and on panels of hundreds of markers even low per-locus error rates accumulate into enough spurious mismatches to inflate Mmin and compromise strict Mmmd duplicate detection. A dedicated SNP pipeline should therefore use mismatch-rate thresholds with explicit error-tolerance bounds on diagnostic panels of dozens to hundreds of markers (Armstrong *et al*. 2025; Flanagan and Jones 2019).

Extending the framework to polyploid accessions and to additional fruit crops is the natural next step, building on the demonstrated feasibility of marker-based pedigree verification in pear (Sawamura *et al*. 2008; Montanari *et al*. 2020). Dedicated SNP and mixed-marker pipelines with rate-based thresholds, network-analysis extensions of the mismatch matrix, and integration with genebank information systems would follow.

## 5. Conclusion

The MMM framework provides deterministic pre-screening of germplasm collections through exact Mendelian exclusion logic, with parameter-free core computation and explicitly reported decision thresholds. Applied to 1,085 apple and 383 sweet cherry accessions from the German Fruit Genebank, MMMAS reproduced published CERVUS parent–offspring assignments with 97–100% concordance, detected one literature-confirmed pair of duplicate genotypes, flagged 20 apple and 10 sweet cherry accessions as genetically distinct, and revealed contrasting MES structures shaped by distinct founder effects and breeding histories.

The Mavg–Mzmp quadrant classification further assigned each apple accession a management-relevant functional role (bridge, pedigree core, genetic extreme, or standard) linking mismatch-based stratification directly to conservation and breeding decisions. Marker-reduction analysis indicated that nine SSR loci retain only the D₀–D₃ core structure (D₄⁺ accessions = 0), that 11 high-PIC loci mark the practical minimum for maintaining diagnostic resolution, and that collections conserving wild relatives or resistance donors require no fewer than 13 loci, with thresholds requiring species- and marker-specific recalibration. Released as open-source software with a bilingual (Chinese/English) interface, MMMAS turns standard SSR fingerprinting data into routine quality-assessment reports for germplasm collections.

## Acknowledgements

We thank the German Fruit Genebank (Julius Kühn-Institut) for making the apple and sweet cherry SSR datasets publicly available in the OpenAgrar repository, and we thank colleagues who provided helpful comments on earlier versions of the manuscript. This work was funded by the National Horticultural Germplasm Repository of China (NHGRC-NH15) and the Ministry of Agriculture and Rural Affairs Germplasm Conservation Project [Project No. 19240429].

## Author contributions

Qiliang Chen conceived the MMM framework, designed the study, performed data analysis and wrote the manuscript. Liuxiu Chen implemented the software and managed code release. Jing Fan and Zi’ang Liu curated the data. Xiaoping Yang and Jingguo Zhang validated the data. Wei Du and Wei Liu reviewed the manuscript. Hongju Hu supervised the project, acquired funding and revised the manuscript. All authors have read and approved the final version.

## Declaration of generative AI and AI-assisted technologies in the writing process

During the preparation of this manuscript (February–September 2026), generative AI tools (Kimi AI, supplemented by ima) were used for English language editing, document formatting, and code development assistance in implementing and refining the MMMAS software. The Mendelian Mismatch Matrix framework, analytical design, validation protocols, and the overall software architecture were conceived by the authors; AI tools supported routine coding and debugging tasks under the authors’ direction. After using these tools, all authors reviewed and tested the code and content, and take full responsibility for the final manuscript.

## Ethical approval

This study used only publicly released microsatellite (SSR) genotypic datasets and published pedigree records of apple (Malus × domestica Borkh.) and sweet cherry (Prunus avium L.) germplasm from the German Fruit Genebank (Julius Kühn-Institut), deposited in the OpenAgrar public database (Broschewitz *et al*. 2023; Höfer *et al*. 2021), together with an additional simulated dataset generated with R code. No new plant material was collected and no genetic resources were accessed, so the Access and Benefit-Sharing (ABS) provisions of the Nagoya Protocol do not apply. The study did not involve human subjects, human tissue, or vertebrate animals, and none of the analyzed datasets contain endangered or protected species. This work complies with the Convention on Biological Diversity (CBD) and the Nagoya Protocol, as well as relevant national and international regulations governing the use of published germplasm data for scientific research.

## Data availability

The apple and sweet cherry SSR genotype data analyzed in this study are publicly available from OpenAgrar (https://doi.org/10.5073/20231220-114634-0 and https://doi.org/10.5073/20210209-092933); pedigree validation data are provided in the supplementary materials of Broschewitz *et al*. (2024) and Reim *et al*. (2023).

## Software availability

Name: MMMAS v1.0.0

License: MIT License

Source code: https://github.com/MMMsystem/MMMAS (tag v1.0.0)

Archived version: https://doi.org/10.5281/zenodo.21441606

Operating system: Windows, macOS, Linux

Dependencies: Python ≥ 3.9, NumPy, pandas; the graphical interface uses the Python standard library (tkinter)

Installation: pip install git+https://github.com/MMMsystem/MMMAS.git, or download from GitHub or Zenodo

Documentation: bilingual README (English/Chinese) included in the repository

Tutorial and example data: provided in the GitHub repository

User support: GitHub Issues

Software copyright: China Software Copyright No. 17864897 (2026SR0650616)

## Abbreviations

MMM: Mendelian Mismatch Matrix
MMMAS: Mendelian Mismatch Matrix Analysis System
Mmn: Mendelian mismatch number
Mmin: Mendelian minimum mismatch number
Mavg: Mendelian average mismatch number
Mzmp: Mendelian zero-mismatch partner number
Mmmd: Mendelian mismatch-mode duplication index
MES: Mendelian Exhaustive Stratification
MGI: Mendelian Grade-gap Index
SSR: simple sequence repeat
SNP: single nucleotide polymorphism
PIC: polymorphism information content
LOD: logarithm of odds

